# Selection on *VPS13A* linked to long-distance migratory behavior in a songbird

**DOI:** 10.1101/585398

**Authors:** David P. L. Toews, Scott A. Taylor, Henry M. Streby, Gunnar R. Kramer, Irby J. Lovette

## Abstract

Animal migration demands an interconnected suite of physiological, behavioral, and neurological adaptations for individuals to successfully navigate and travel over long distances^1–3^. This trait complex is especially crucial for small songbirds whose migratory behaviors—like directionality and orientation—are innate^4,5^, rather than being learned as in many large, longer-lived birds. Identifying causal genes involved in these traits has been a central goal of migration ecology, and this endeavor has been furthered by genome-scale comparisons^6^. However, even the most successful studies of migration genetics have only achieved low resolution associations, identifying large chromosomal regions, across multiple haplotype blocks, that encompass hundreds of putatively causal genes^7,8^. Here we leverage the extreme genomic similarity among golden-winged (*Vermivora chrysoptera*) and blue-winged warblers (*Vermivora cyanoptera*)^9^ to identify a single gene—*Vacuolar Protein Sorting 13 Homolog A* (*VPS13A*)—that is associated with distinct differences in migration directionality to Central American (CA) versus South American (SA) wintering areas^10^. Moreover, we find significantly reduced sequence variation in this gene-region for SA wintering birds, and show this is the result of strong natural selection on this gene. In humans, *VPS13A* codes for chorein, and variants of this gene are associated with the neurodegenerative disorder chorea-acanthocytosis^11^. This new association provides the strongest gene-level linkage for avian migration directionality, and further interrogation of this gene will allow for a better understanding of its role in neuro-muscular processes across vertebrates.

---

All *Vermivora* warblers breed in North American and migrate annually to over-winter in two disjunct Neotropical regions: in South America (SA), near the Venezuelan and Columbian border, and in Central America (CA), from Northern Panama to Guatemala^10,12,13^. These overwintering locations are highly predictable based on breeding locations and plumage phenotype: golden-winged warblers breeding in the Appalachian Mountains winter in SA, whereas golden-winged warblers breeding outside of Appalachia—as well as most blue-winged warblers—winter in CA^10^.

Golden-winged and blue-winged warblers represent a system with exceptional utility for discovering possible gene-migration associations. This is because much of the genetic variation amongst these warblers is shared—even between individuals differing strongly in their plumage phenotypes^9^—due to a long history of gene flow. This unusually homogeneous genomic background greatly enhances our power to identify gene-level trait associations—such as those linked to different migration behaviors—compared to typical inter-species comparisons in which much higher background divergence and population structure confound genotype-phenotype associations.

To test for associations between SNPs and wintering location, we used low-coverage whole-genome resequencing from 70 *Vermivora* warblers classified as either wintering in CA or SA. These samples either had full geolocator tracks associated with them to determine overwintering location (*n* = 44)^13^, or were reliably classified based on breeding location and plumage type (*n* = 26). We aligned reads (mean = 30 million reads / individual) to the Myrtle Warbler reference genome^9^ and assayed variation across the resulting 8.3 million variable SNP sites (mean depth per SNP = 4.9 reads).

We found many clustered SNPs associated with overwintering behavior within a 120kb region of warbler scaffold 24, a region that aligns to the Zebra Finch Z chromosome (Figure 2). There is no single SNP with a highly significant association (*i.e.*, −log_10_(*P*) > 7) in this small region. However, there are an exceptionally high number of moderately significant SNPs (16% of 349 SNPs with −log_10_(*P*) > 5): in one 10kb window within this region there are 51 SNPs with −log_10_(*P*) > 5. By contrast, the next highest window in the entire genome has only 7 such SNPs and the mean number of SNPs per 10kb window with a −log_10_(*P*) > 5 across the *Vermivora* genome is 0.004.

**Figure 1 –.**
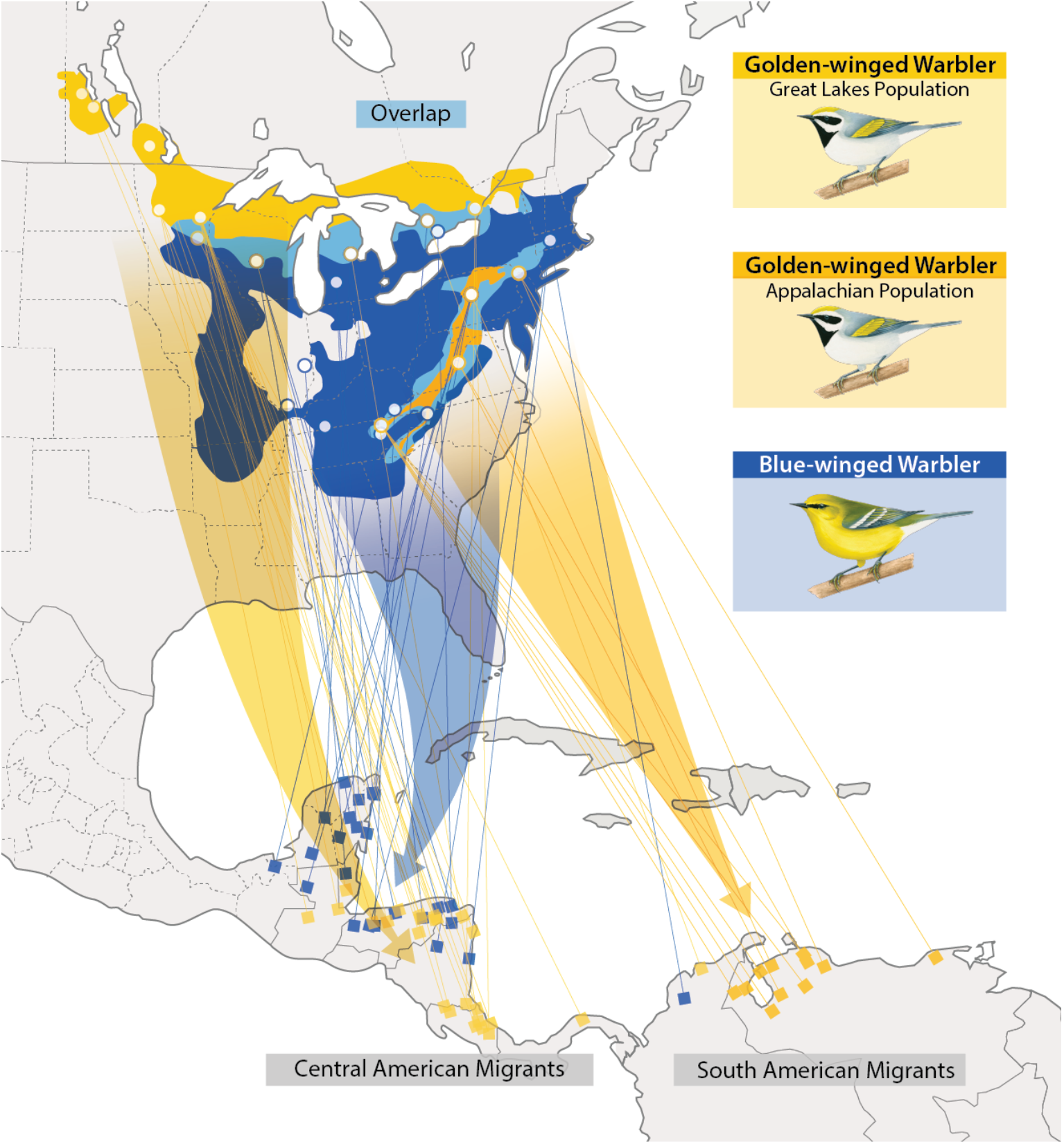
Distribution and migration patterns of *Vermivora* warblers from geolocator data. Blue-winged warblers (blue) breed at lower elevations and lower latitudes, and primarily migrate to and from Central America (CA). There are two breeding populations of golden-winged warblers: the Great Lakes population (lighter orange) also migrate to and from Central America, whereas the Appalachian population migrates to South America (SA; darker orange). Circles represent geolocator deployment location during the breeding season; squares represent predicted non-breeding locations. Migration data and maps adapted from [10] by Jillian Ditner. Illustrations by Liz Clayton Fuller.

**Figure 2 –.**
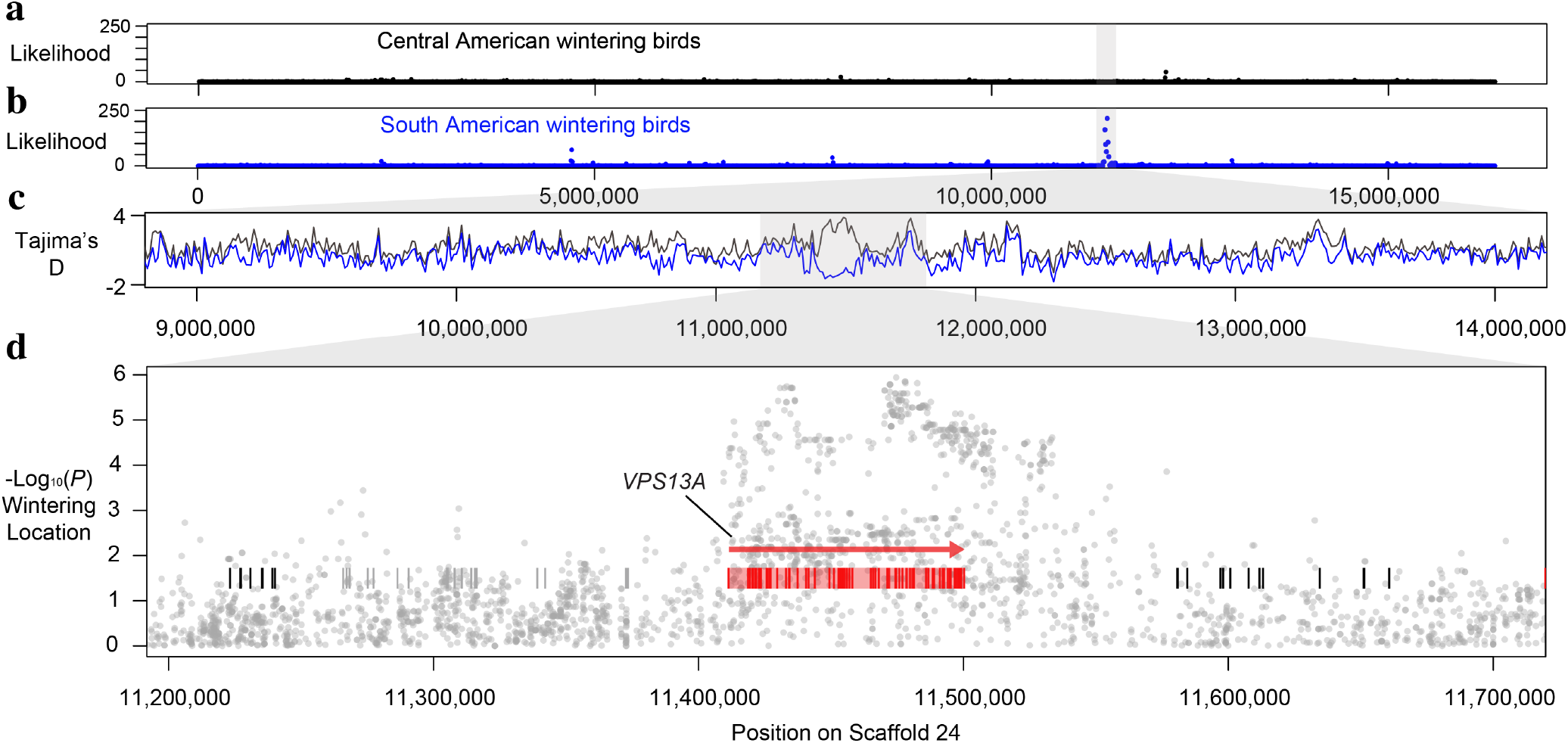
Patterns of genetic variation and associations with *Vermivora* wintering locations. **Panels (a)** and **(b)** show the composite likelihood ratio tests of a selective sweep—from SweeD^14^—for birds wintering in South America (blue) or in Central America (black) for a portion of the Z chromosome (warbler scaffold 24 shown). **Panel (c)** shows estimates of Tajima’s D for a smaller subset of this scaffold. The region outlined in grey is the only portion of the entire genome that shows highly different patterns between the groups for Tajima’s D (Figure 4). **Panel (d)** shows the statistical association between SNP genotypes and overwintering location for this portion of the genome. While no single SNP shows a highly statistically significant association (*i.e.*, −log_10_*P* > 7), the clustering of moderately significant SNPs in this region is an extreme outlier compared to the rest of the genome. A single gene—*VPS13A* (red)—falls within this region of association.

This region of the Z chromosome is also an outlier in allele frequencies between CA and SA birds (Figure 3): the average genome-wide *F*_ST_ between CA and SA wintering birds is 0.002. In this region, however, *F*_ST_ is 0.22, which is two orders of magnitude higher than the genome-wide average. Moreover, the top 12 most differentiated 10kb *F*_ST_ windows across the *Vermivora* genome fall within this region. The association and *F*_ST_ results together imply that: 1) variation within this region of the Z chromosome is an outlier in comparisons of SA versus CA birds; and 2) that these differences are not driven by fixed SNPs, but rather the variation within SA individuals in this region is a subset of the variation present in CA wintering birds.

**Figure 3 –.**
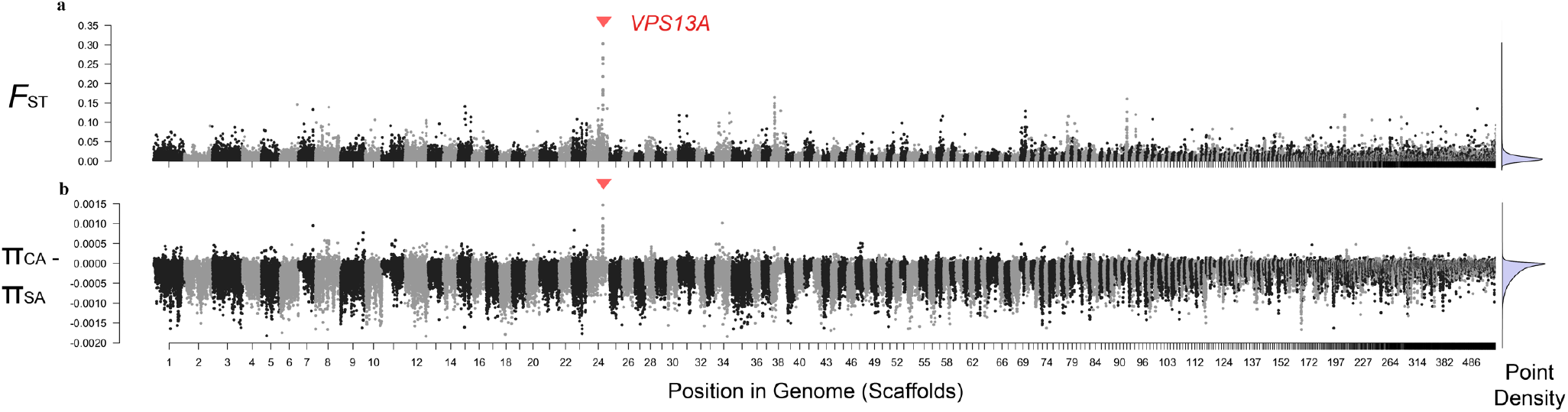
Patterns of genome-wide differentiation and diversity between South American (SA) and Central American (CA) wintering *Vermivora* warblers (both with a mix of golden-winged and blue-winged phenotypes). The region around *VPS13A* is an outlier between birds wintering in different areas, and variation within SA is a significantly reduced subset of what is found in CA birds. **Panel (a)** shows estimates of *F*_ST_ for 10kb non-overlapping windows across the genome (with at least 4 variants per window). **Panel (b)** shows the *difference* in nucleotide diversity (π) between CA and SA wintering birds. Across the genome, Δπ is highly comparable between the two groups (*i.e.*, mean of the difference is ~0). However, the region containing *VPS13A* shows a 42% reduction in diversity in SA wintering birds. Given the large number of overlapping points, kernel densities of points are shown in the right panels.

This pattern has likely been produced by strong natural selection (Figure 3). We additionally found that the divergent region associated with overwintering location has reduced DNA sequence variation in SA birds compared to those wintering in CA. Comparing nucleotide diversity between SA and CA wintering individuals, there is consistently very little difference between the groups across the genome: the average genome-wide difference in diversity in 10kb windows (*i.e.*, π_CA_ − π_SA_) is −0.0002. By contrast, the candidate region linked to wintering differences shows a 42% reduction in diversity for SA birds compared to CA birds (mean π_SA_ = 0.0007; mean π_CA_ = 0.001), with no other genomic region showing such a large difference in diversity (Figure 3b). Tajima’s D (TajD)—a statistic that helps to distinguish between DNA sequences evolving neutrally versus under non-neutral processes—also implies this candidate region is a genomic outlier (Figure 4). Genome-wide, TajD is highly correlated between SA and CA wintering birds (Pearson’s correlation = 0.85), with CA birds having consistently higher values compared to SA birds across their genomes. Reduced TajD in SA birds is likely a result of persistent population declines in Appalachian golden-winged warblers, which has been associated with wintering-site factors^10^. TajD estimates for windows within the region of the Z chromosome associated with migratory behavior, however, have values much lower than the genome-wide average (red points in Figure 4). We also tested for natural selection explicitly with SweeD^14^ (sweep detector) and found a very high likelihood of a positive selective sweep in SA birds: the composite likelihood ratio (CLR) for SA birds in the migration-associated region is 213; CLR for this same region in CA wintering birds is 0.099. Taken together, the various metrics and analyses (π, TajD, and SweeD) imply reduced diversity due to strong selection in SA birds.

**Figure 4 –.**
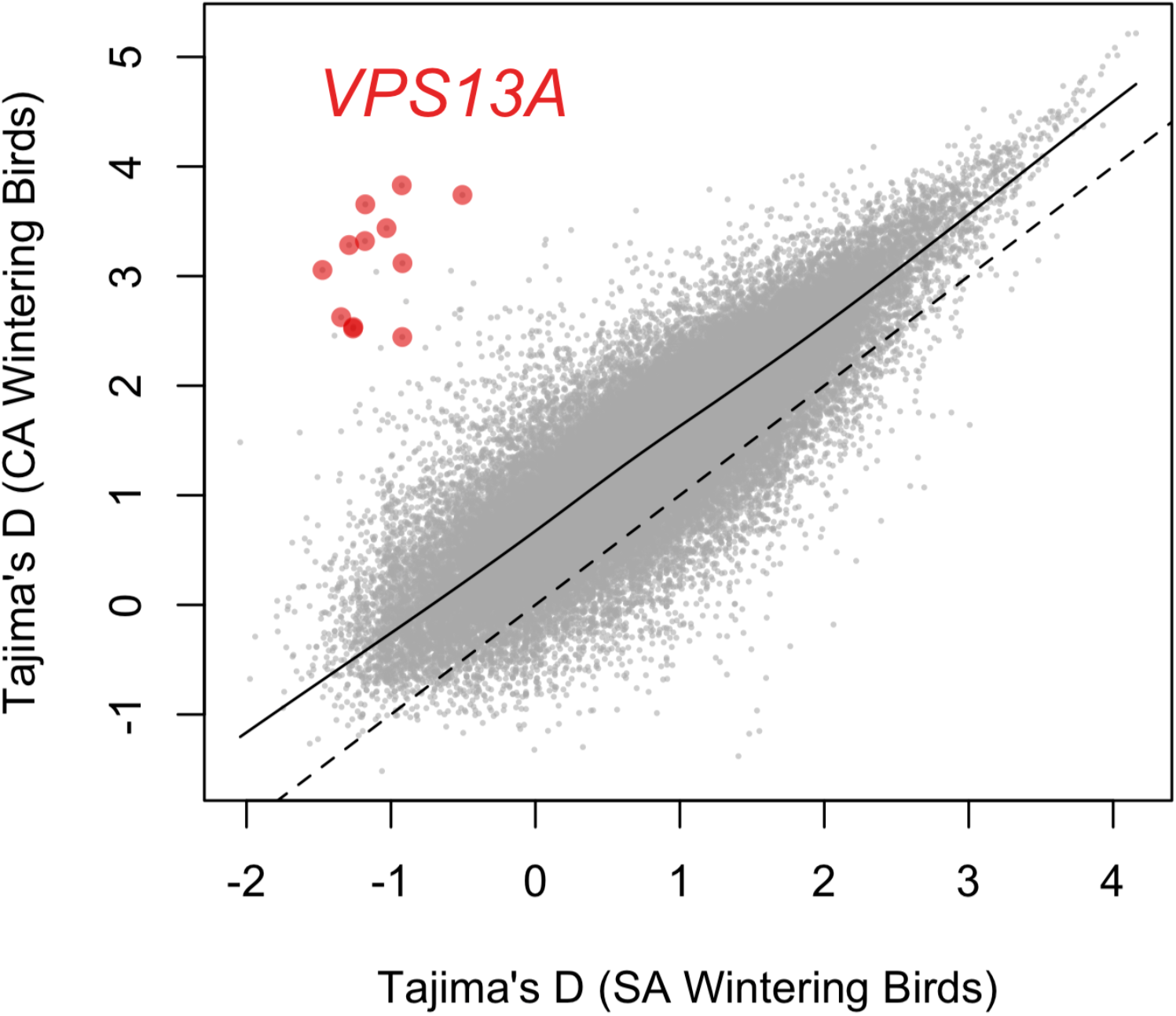
Estimates of Tajima’s D for 110,317 non-overlapping 10kb windows for South American wintering birds and Central American wintering birds. Overall, there is a positive correlation between the groups across the genome, although CA wintering birds have higher values on average (the dotted line shows a 1:1 relationship). This reduction in Tajima’s D in SA wintering birds is possibly due to population declines, likely associated with wintering-site factors^10^. The 12 windows that fall within the region on the Z chromosome associated with wintering location, and include the gene *VPS13A*, do not follow this genome-wide pattern: these windows have very low Tajima’s D values in SA birds, compared to much higher values for those same windows in CA wintering birds.

Only one gene—*VPS13A*—falls within the region associated with wintering location in *Vermivora* warblers. Comparing CA and SA wintering birds, there are 12 SNPs with *F*_ST_ >0.3 that fall within the exons of *VPS13A*. Of these, only one—in exon 45—has a non-synonymous substitution: G-1792^Ser^ is at high frequency in SA birds, whereas C-1792^Thr^ is common in CA wintering birds (amino acid locations are relative to NCBI *Serinus canaria* assembly; Accession Number GCA_000534875.1). For all SNPs within exons, the exon 45 SNP in *VPS13A* has the highest *F*_ST_ (0.41) between CA and SA birds. This SNP is also <500 bp from an intronic SNP that is the most differentiated between SA and CA individuals across the entire genome (*F*_ST_ =0.57). SA birds have retained the ancestral state (G-1792^Ser^) at the differentiated non-synonymous exon SNP, whereas CA individuals have the derived allele. Across all sequenced avian genomes (Zebra finch, taeGut3.2.4; Collared Flycatcher, FicAlb_1.4; and Chicken, GRCg6a), as well as the green anole genome (AnoCar2.0), this “G” variant is conserved, consistent with a possible functional role for this amino acid substitution.

*VPS13A* does not have a characterized function in birds. However, it has a well-known disease phenotype in humans: *VPS13A* codes for chorein, and variants of this gene are associated with the neurodegenerative disorder chorea-acanthocytosis^15^. This disease is inherited as an autosomal recessive condition that includes movement disorders, chorea, and dystonia^16^. Recent work also shows *VPS13A* as closely associated with mitochondria, and may be involved in lysosomal degradation^17^. Whereas the phenotypes associated with *VPS13A* present an intuitive connection to the neurological and energetic demands of migration, the role of the *VPS13A* protein in specific molecular processes in most organisms is still unclear, and thus its exact role in avian migration remains speculative.

Determining how specific mutations in *VPS13A* are associated with migratory phenotypes, understanding how this gene relates to specific variation in migratory directionality, and quantifying the specific mechanism(s) of selection acting in SA wintering birds, should all be the target of future research in migratory *Vermivora* warblers. The finding that variation in *VPS13A* is both linked to migratory differences in *Vermivora* warblers, and has experienced strong natural selection, also makes it an excellent candidate for further study in other migratory taxa.

## Supporting information

Supplemental Table 1, sample information

## METHODS

### Sampling

We used blood samples, taken from the brachial vein, from breeding warblers captured between 2008 and 2017 (*n* = 70). These were from individuals either previously wintered in Central America (CA; *n* = 25 golden-winged phenotypes, *n* = 23 blue-winged phenotypes, and *n* = 2 hybrid phenotypes) or South America (SA; *n* = 17 golden-winged phenotypes, *n* = 1 blue-winged phenotype, *n* = 2 hybrid phenotypes). Most individuals were territorial males captured during the breeding season, although we also included 4 samples of breeding females. For 44 individuals, we obtained full migration track data from light-level geolocators. These data have been presented previously^10,13^. Briefly, geolocators are simple, lightweight tags that record levels of ambient light at regular intervals, enabling the estimation of location based on seasonal variation in spatio-temporal patterns in sunrise and sunset.

We captured adult *Vermivora* warblers at sites across their breeding distribution and outfitted them with geolocators that we retrieved during the following breeding season (*i.e.*, following one annual migration cycle). We analyzed geolocator data to derive spatially explicit probability density functions for individual warblers during the winter period^13,18^. We classified the wintering location of individual warblers based on the location of the maximum likelihood cell (0.5°) of the winter probability density function (*i.e.*, occurring in CA or SA). Across all geolocator samples, individuals were unambiguously assigned to CA or SA wintering sites. For those individuals where we did not have geolocator data (*n* = 26), we included only those samples from sites where there is no evidence of broad-scale variation in overwintering location and thus could be predicted unambiguously (*i.e.* only broadly allopatric golden-winged and blue-winged warblers, which winter in CA, and Appalachian golden-winged warblers, which winter in SA).

### Genome library preparation and filtering

Our genomic analysis relied on low-to-moderate coverage whole genome resequencing data. We divided our sequencing arbitrarily into four batches. We extracted DNA from all samples using UPrep spin columns (Genesee). For one sequencing batch, we used the Illumina TruSeq PCRFree kit, following the protocols to generate libraries with a 350 bp insert. For the three other batches, DNA concentrations were lower, and therefore we used the Illumina TruSeq Nano kit—which includes an 8-cycle PCR enrichment—also producing 350 bp insert sizes. We sequenced these libraries across four lanes of an Illumina NextSeq using the paired-end 150 bp sequencing chemistry. In some cases, we combined *Vermivora* samples with other wood warblers from other projects, but consistently included 24 individuals per sequencing lane.

To remove adapters, trim low quality bases from the ends of reads, and collapse overlapping pairs, we used AdapterRemoval 2.1.1 with the -collapse -trimns -minlength 20 -qualitybase 33 options. We then used the very-sensitive-local set of presets in Bowtie2 to align reads to the myrtle warbler genome ^9^. For those samples where PCR enrichment was used, we used the MarkDuplicates command in Picard Tools 2.8.2 to identify and mask likely PCR duplicates.

To call SNPs we used the UnifiedGenotyper in GATK 3.8 and the VariantFiltration function with the expression ““QD<2.0, FS>40.0, MQ<30.0, HaplotypeScore>12.0, MappingQualityRankSum<−12.5, ReadPosRankSum<−8.0”. Using VCFTools, we filtered genotypes with a minimum genotype quality (minGQ) <8 as missing data. We used a minimum allele frequency of 5% and removed loci with more than 30% missing data.

### Data analysis

We used the genome-wide efficient mixed model association algorithm (GEMMA 0.98.1^19,20^) to associate variation at SNPs with the binary classifications of overwintering location. GEMMA requires complete or imputed SNP data. Therefore, we first used BEAGLE (v. 4.1^21^) to impute missing data for those sites with missing data. We transformed our data into binary BED format using PLINK 1.9^22^. We used the Wald test to identify significant associations between SNP genotypes and wintering location and report the p-values using a univariate linear mixed models. To visualize the resulting associations, we used a modified version of the ‘qqman’ package in R to generate Manhattan plots^23^. We estimated *F*_ST_, π, and Tajima’s D using unimputed genotypes in VCFtools^24^. We conducted a scan for positive selection using SweeD on separate VCF files for SA versus CA wintering birds. This program uses the site frequency spectrum to calculate the composite likelihood ratio (CLR) of a selective sweep model over a neutral model. We conducted this test for 10kb windows across warbler scaffold 24.

## ACKNOWLEDGEMENTS

The authors would like to thank D. Andersen, D. Buehler, and P. Wood, co-PIs with H. Streby on the *Vermivora* geolocator project. The authors would also like to thank Steve VanWilgenburg, Jeff Larkin, Rachel Vallender, Robert Ricklefs, and Kyle Aldinger for helping collect and contribute blood samples. Jillian Ditner and Liz Clayton Fuller produced the illustrations.

## AUTHOR CONTRIBUTIONS

DPLT and ST designed the study. DPLT performed the analyses and wrote the original draft of the manuscript. HS and GK analyzed the migration track data. All authors contributed to the editing of the manuscript.

